# An *Escherichia coli* Chassis for Production of Electrically Conductive Protein Nanowires

**DOI:** 10.1101/856302

**Authors:** Toshiyuki Ueki, David J.F. Walker, Trevor L. Woodard, Kelly P. Nevin, Stephen S. Nonnenmann, Derek R. Lovley

**Author notes:** Authors contributed equally (Toshiyuki Ueki, David J.F. Walker, Derek R. Lovley).

## Abstract

*Geobacter sulfurreducens’* pilin-based electrically conductive protein nanowires (e-PNs) are a revolutionary electronic material. They offer novel options for electronic sensing applications and have the remarkable ability to harvest electrical energy from atmospheric humidity. However, technical constraints limit mass cultivation and genetic manipulation of *G. sulfurreducens*. Therefore, we designed a strain of *Escherichia coli* to express e-PNs by introducing a plasmid that contained an inducible operon with *E. coli* genes for type IV pili biogenesis machinery and a synthetic gene designed to yield a peptide monomer that could be assembled into e-PNs. The e-PNs expressed in *E. coli*, and harvested with a simple filtration method, had the same diameter (3 nm) and conductance as e-PNs expressed in *G. sulfurreducens*. These results, coupled with the robustness of *E. coli* for mass cultivation and the extensive *E. coli* toolbox for genetic manipulation, greatly expands opportunities for large-scale fabrication of novel e-PNs.

## Introduction

Electrically conductive protein nanowires (e-PNs) show promise as revolutionary, sustainably produced, and robust electronic materials ^1-8^. They are biocompatible and can readily be adapted for a multitude of sensing applications ^1-7^. Devices comprised of thin layers of e-PNs function as ‘humidity-powered electrical generators’, continuously harvesting energy in the form of electricity from atmospheric humidity ^8^. However, the implementation of e-PNs in electronic devices has been limited due to a lack of methods for mass production. *In vitro* assembly of peptides into conductive nanofilaments is feasible ^9, 10^, but the filaments tend to agglomerate into gels at high peptide concentrations, limiting possibilities for large-scale fabrication. Furthermore, synthesis of the peptide monomers required for *in vitro* assembly is expensive, potentially limiting e-PN affordability.

*In vivo* assembly of e-PNs with microorganisms has several advantages over *in vitro* synthesis. Benefits include much lower cost and greater flexibility in e-PN design options with a production platform fueled with inexpensive, renewable feedstocks. A diversity of bacteria and archaea assemble peptides that show homology to bacterial type IV pilins into e-PNs ^11-15^, but the e-PNs of *Geobacter sulfurreducens* have been most intensively investigated ^1, 16^. *G. sulfurreducens* e-PNs can be fabricated *in vivo* with acetate as the carbon and energy source. Once the cells are grown, the e-PNs can be harvested, retaining their conductive properties ^12, 17-19^.

The exquisite machinery that bacteria possess to assemble pilin proteins into filaments ^20^ confers great control over e-PN production, yielding a highly uniform product. The microbial assembly process also offers substantial opportunities for producing diverse, new types of e-PNs. For example, the conductivity of e-PNs produced with *G. sulfurreducens* has been tuned over a million-fold with simple modifications to the *G. sulfurreducens* pilin gene to either increase or decrease the abundance of aromatic amino acids ^1, 16^. Pilin genes can be designed to encode additional peptides at the carboxyl end of the pilin, yielding e-PNs that retain their conductivity and display the added peptides on the outer surface of the wires ^7^. This peptide display along the wires offers unique possibilities for introducing peptide ligands to confer specific sensing functions to e-PN devices with a flexibility in sensor design not feasible with other materials such as carbon nanotubes or silicon nanowires ^7^. Peptides displayed on the outer surface of e-PNs might also be designed to promote e-PN binding to surfaces to facilitate wire alignment or to function as chemical linkers with polymers for the fabrication of composite materials ^7^. Synthetic gene circuits introduced to control the expression of multiple e-PN monomer genes within a single cell offer the possibility to further tune e-PN function by producing heterogeneous wires comprised of multiple types of e-PN monomers with the stoichiometry of each monomer type precisely controlled ^7^. These design options would be difficult to replicate with *in vitro* assembly of e-PNs or fabrication of nanowires from non-biological materials.

*Geobacter*-fabricated e-PNs have several other advantages over traditional non-biological nanowire materials. Production of the e-PNs requires 100-fold less energy than is required for fabricating silicon nanowires or carbon nanotubes ^2^. No toxic chemicals are required for e-PN fabrication and the final product is biocompatible, environmentally benign, and recyclable ^2^. Yet, e-PNs are remarkably robust, maintaining function even under harsh CMOS-compatible fabrication conditions ^6^. Proof-of-concept studies have demonstrated the dynamic sensing response of *Geobacter*-fabricated e-PNs; the ability of these e-PNs to function as the conductive component in flexible electronics; and that, in the appropriate electrode/e-PN configurations, thin films of *Geobacter* e-PNs can generate electricity from the humidity naturally present in air ^6-8, 18^.

Barriers to large-scale e-PN production have been a limitation to realizing the potential of *Geobacter* e-PNs for these and other possible applications. *G. sulfurreducens* must be grown anaerobically to produce e-PNs. This requirement adds technical complexity and costs. A strain of *Pseduomonas aeruginosa*, grown aerobically, heterologously expressed the *G. sulfurreducens* pilin gene with the assembly of e-PNs with properties similar to the e-PNs expressed in *G. sulfurreducens* ^21^. However, *P. aeruginosa* is a pathogenic microorganism and thus not ideal for large-scale commercial production of e-PNs. Furthermore, the expression of the e-PNs in *P. aeruginosa* remained under the control of the native regulatory system, limiting options for controlling the timing and extent of e-PN expression ^21^.

We thought that *Escherichia coli* might be an ideal chassis for e-PN fabrication. *E. coli* is a common platform for the commercial scale production of organic commodities ^22, 23^. The substantial *E. coli* genetic toolbox, including the possibility of introducing unnatural amino acids ^24, 25^, could provide broad options for designing e-PNs with different properties and functions. Non-pathogenic strains of *E. coli* typically do not express type IV pili. However, introduction of an artificial operon of pilus assembly protein genes from pathogenic *E. coli* into non-pathogenic *E. coli* yielded a non-pathogenic strain that expressed the same type IV pili that pathogenic *E. coli* express ^26^. This finding, and the fact that bacteria will often assemble heterologous pilins into pili ^12, 13, 21, 27-29^, suggested that it might be possible to develop a non-pathogenic strain of *E. coli* that would express *Geobacter* e-PNs.

Another limitation to large-scale *in vivo* production of e-PNs has been the methods for separating the e-PNs from cells. Previously described methods have included multiple laborious steps, often with strategies such as ultracentrifugation and/or salt precipitation procedures that would be difficult to economically scale ^12, 17^.

Here we report on the construction of a strain of *E. coli* amended with genes for the expression of type IV pili assembly machinery and a synthetic gene designed to yield e-PNs comparable to those produced by *G. sulfurreducens*. This strain produces e-PNs with characteristics similar to the e-PNs expressed by *G. sulfurreducens*. Simple aerobic growth of the *E. coli* designed for e-PN production, coupled with a new simplified method for harvesting e-PNs from cells, suggests that large-scale production of e-PNs will be feasible.

## Results and Discussion

We engineered a standard lab strain of *E. coli* to produce e-PNs by introducing the genes known from previous studies ^26^ to be required for type IV pili biogenesis. We modified the previous design of *E. coli* ^26^, which was constructed to produce pili from the *E. coli* pilin protein PpdD, as follows: 1) the gene for pilin monomer is within a separate cloning site to make it convenient to exchange the gene for the pilin of interest (Figure 1a); 2) the gene clusters were constructed with common and less expensive restriction enzymes to aid in modularity (Figure 1b); 3) ribosome binding sites for *hofB, hofM*, and *ppdA* were changed to improve translation efficiency (Figure 1c); 4) a gene cluster containing *ppdA, ppdB, ygdB*, and *ppdC* and *gspO* were connected by 2-step PCR instead of using a restriction enzyme to delete unnecessary sequence (Figure 1b,d); 5) intergenic regions were shortened to delete unnecessary sequence (Figure 1d); 6) the *tac* promoter ^30^, one of the strongest promoters in *E. coli*, was incorporated to enhance transcription of the genes for the assembly of type IV pili (Figure 1b); and 7) a gene for the LacI repressor was included in the expression vector to repress the genes when desired, such as during cloning (Figure 1a). We also deleted the gene *fimA* to prevent the formation of type I pili using previously described genetic methods ^31, 32^.

**Figure 1.**
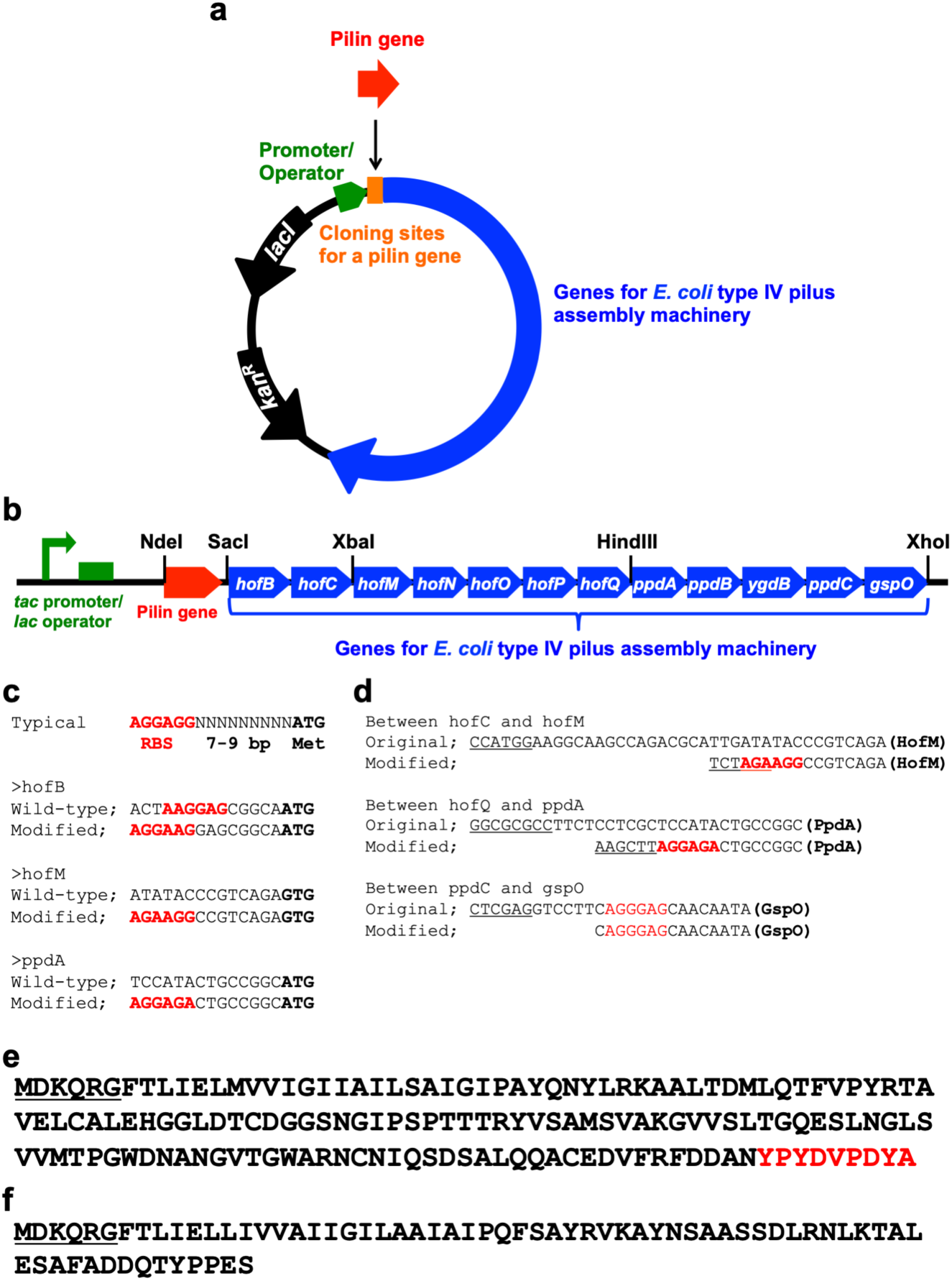
Construction of type IV pilus assembly system in *E. coli* and synthetic peptide monomer for e-PN assembly. (a) Expression vector for genes for type IV pilus assembly in *E. coli*. (*lacI*, Lac repressor gene; *kan*^*R*^, kanamycin resistance gene); (b) Gene organization for *E. coli* type IV pilus assembly with cloning sites designated; (c) Ribosome binding sites (designated in red) that were changed from those in previous studies ^26^; (d) Intergenic regions in the synthetic operon that were changed from those in previous studies ^26^ (ribosome binding sites in red, restriction enzyme sites underlined); (d) Amino acid sequence of the HA-tagged PpdD (signal sequence underlined, HA tag sequence in red); (f) Amino acid sequence of the synthetic peptide monomer designed for assembly in e-PNs, which is a combination of a portion of the *G. sulfurreducens* pilin, PilA, with the *E. coli* PpdD signal sequence (underlined).

The initial strain was constructed with a pilin gene that added the HA tag (YPYDVPDYA) at the carboxyl terminal end of the *E. coli* pilin protein PpdD (Figure 1e). With this modification the tagged pilin protein (PpdD-HA) could readily be detected with the commercially available antibody for the HA tag. PpdD-HA was detected in the cell extract from the strain containing the genes for PpdD-HA pilus assembly but not in extracts from a control strain that lacked the gene for PpdD-HA (Figure 2a). PpdD-HA pili were sheared from cells and PpdD-HA was detected in the sheared fraction from the strain containing the genes for PpdD-HA pilus assembly, but not from the control strain without the PpdD-HA gene (Figure 2a). These results confirmed that the modified expression system for type IV pilus assembly was effective for pili production.

**Figure 2.**
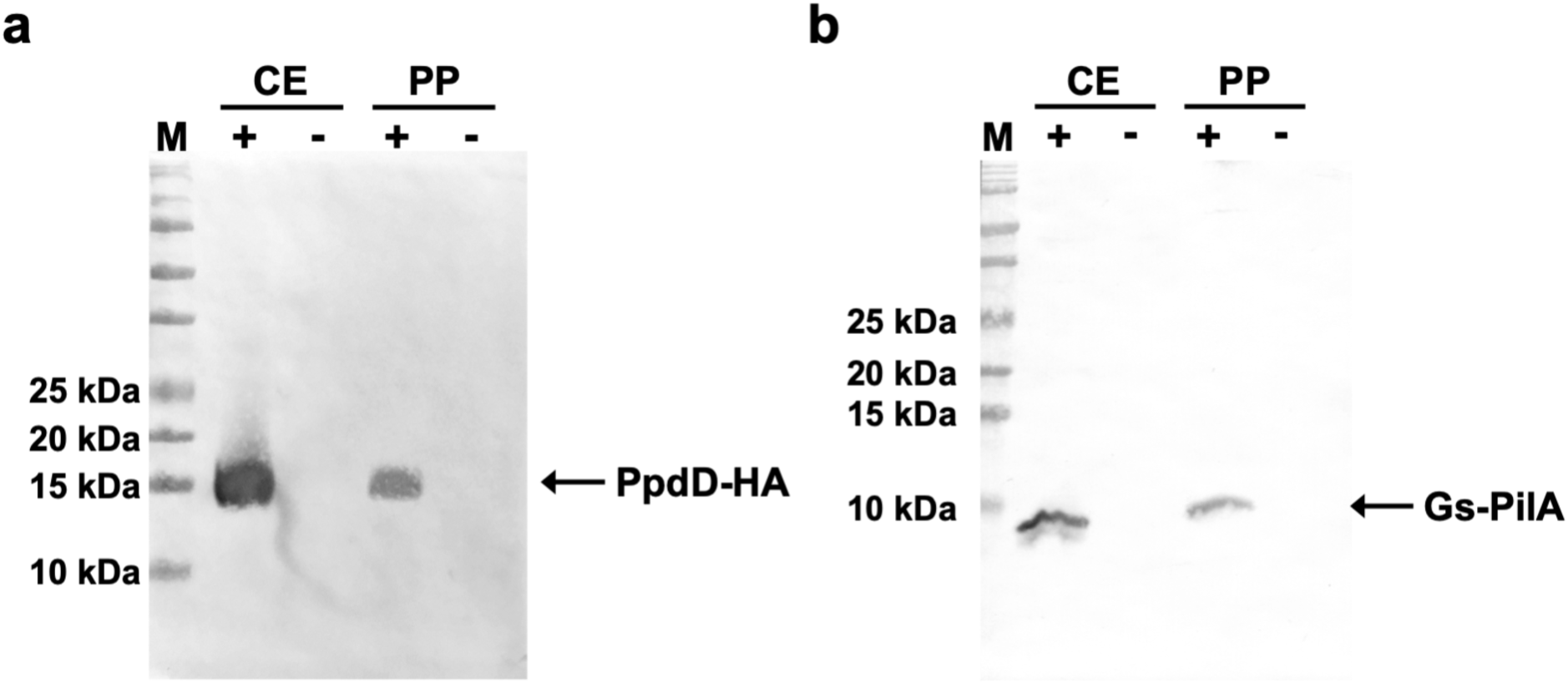
Expression of pili and e-PNs in *E. coli*. (a) Results with the strain of *E. coli* with genes for pilus assembly and the gene for PpdD-HA. (b) Results from *E. coli* strain GPN, which contained genes for pilus assembly and the gene for a synthetic pilin monomer designed to yield e-PNs assembled from a modified *G. sulfurreducens* PilA monomer. Western blot analyses of denatured proteins separated on an SDS PAGE gel and analyzed with antibody for (a) the HA tag on PpdD-HA pilin or (b) *G. sulfurreducens* PilA. Strains with the pilin genes designated with (+). Control strains without the pilin genes designated (-). Samples from whole cell extracts (CE) and the pili preparations (PP) were examined. Lanes designated M show molecular weight standard markers.

To express e-PNs similar to those expressed in *G. sulfurreducens* in *E. coli*, we designed a gene to yield a synthetic peptide monomer for assembly into e-PNs. The peptide was similar to the *G. sulfurreducens* pilin monomer, PilA, with the exception that the signal sequence was replaced with the *E. coli* PpdD signal sequence to facilitate e-PN assembly in *E. coli* (Figure 1f). The gene for the synthetic e-PN monomer was cloned into the location designated ‘pilin gene’ (Figure 1a). The strain with the synthetic gene for the e-PN monomer was designated *E. coli* strain GPN (*<u>G</u>eobacter* <u>p</u>rotein <u>n</u>anowire). The e-PN monomer was detected in whole cell extracts of strain GPN with PilA antibody, but not in the control strain that lacked the gene for the e-PN monomer (Figure 2b).

e-PNs were harvested from strain GPN with physical shearing from the cells, as in previous studies of e-PNs expressed in *G. sulfurreducens* ^17^. In those previous studies, the cells were separated from the sheared e-PNs with centrifugation and then the e-PNs in the supernatant were collected with ultracentrifugation or ammonium sulfate precipitation ^17^. These methods of e-PN collection are labor intensive and will be difficult to adapt to large-scale production. Therefore, the e-PNs sheared from strain GPN and separated from cells were treated with Triton X100 detergent and then were collected on a filter with a 100 kDa molecular weight cutoff. This method is simpler and faster than previously described ^17^ e-PN purification methods.

The e-PNs harvested from *E. coli* strain GPN were ca. 3 nm in diameter and several µm in length (Figure 3a), a morphology similar to the e-PNs expressed in *G. sulfurreducens*. No filaments were observed in similar preparations when the gene for the e-PN monomer was omitted from *E. coli* strain GPN. Denaturation of the e-PNs from *E. coli* strain GPN yielded a monomer that reacted with PilA antibody whereas the monomer was not detected in preparations from the control strain without the gene for the e-PN monomer (Figure 2b).

**Figure 3.**
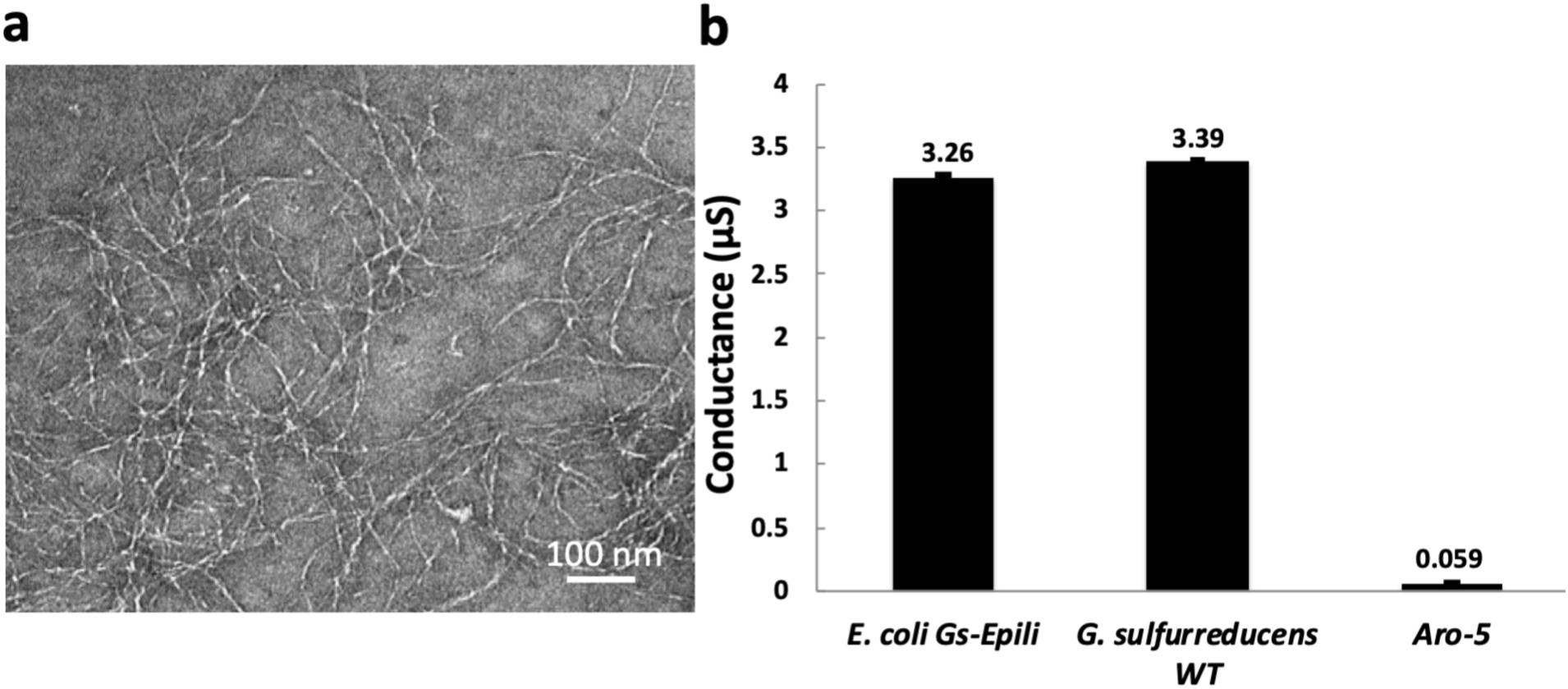
Characterization of the e-PNs expressed in *E. coli* strain GPN. (a) Transmission electron micrograph. (b) Conductance of films of e-PNs expressed in *E. coli* strain GPN compared with e-PNs from wild-type *Geobacter sulfurreducens* and the Aro-5 strain of *G. sulfurreducens*. The results are the means and standard deviation of triplicate measurements on each nanoelectrode array, and at least three independent nanoelectrode arrays. Results for wild-type *G. sulfurreducens* and strain Aro-5 were published previously ^13^.

The conductance of thin films of the e-PNs from *E. coli* strain GPN, determined with a nanoelectrode array as previously described ^13^, was 3.26 ± 0.35 μS, similar to the conductance of 3.39 ± 0.04 μS for e-PNs harvested from *G. sulfurreducens* (Figure 3b). The conductance of these e-PNs was much higher than the conductance of wires harvested from strain Aro-5 (Figure 3b), a strain of *G. sulfurreducens* that expresses a synthetic pilin gene designed to yield protein nanowires with low conductivity ^18, 27^.

The conductance of individual e-PNs was evaluated on highly oriented pyrolytic graphite with atomic force microscopy employing a conductive tip, as previously described ^15^. The diameter of the e-PNs was 3 ± 0.04 nm (n=18; 6 measurements on 3 independent pili) the same as e-PNs expressed with *G. sulfurreducens* (Figure 4). The e-PNs were conductive with an ohmic-like current-voltage response (Figure 4d). The conductance of the individual e-PNs, 4.3 + 0.8 nS (n=9), compared well with the previously reported ^15^ conductance of 4.5 ± 0.3 nS for individual e-PNs expressed in *G. sulfurreducens*.

**Figure 4.**
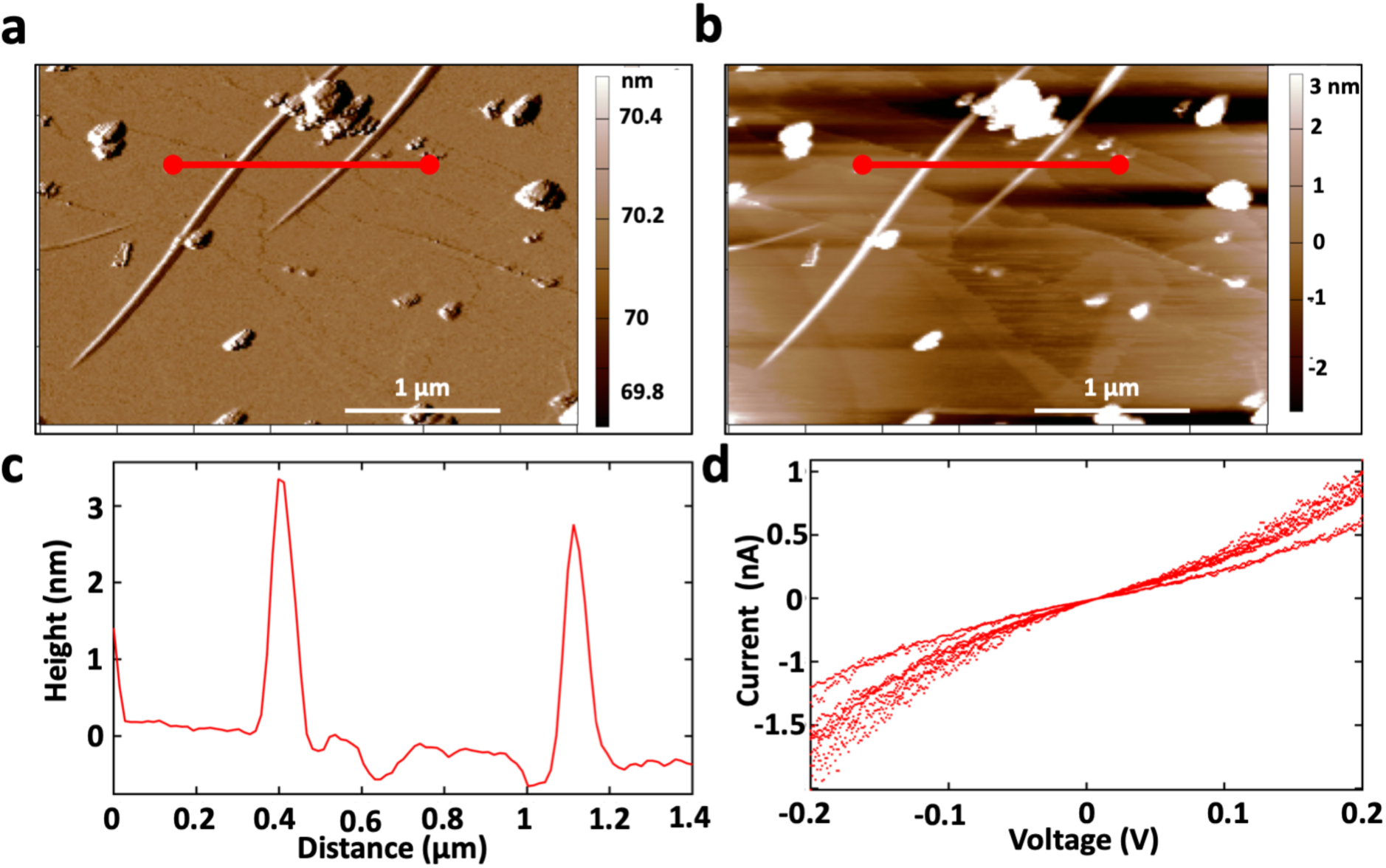
Characterization of individual e-PNs expressed in *E. coli* strain GPN. (a) Amplitude image of 2 e-PNs in amplitude modulation mode (b) Height image of each e-PN (c) Cross-section line trace showing the height of 2 individual e-PNs, designated in panel a and b by the redline, demonstrating ∼3 nm height (diameter). (d) Current voltage response of nine individual measurements (3 measurements on 3 e-PNs).

## Conclusions

The fabrication of e-PNs with *E. coli* offers substantial advantages over expression in *G. sulfurreducens*. Special equipment and expertise is required to anaerobically culture *G. sulfurreducens*, whereas *E. coli* can be simply grown under ambient, aerobic atmospheric conditions. When coupled with our finding that the e-PNs can be collected with simple filtration method, expression of e-PNs in *E. coli* offers the potential for large-scale e-PN production.

Furthermore, e-PN expression in *E. coli* offers much greater flexibility for the design of a wider diversity of e-PNs than would currently be possible with *G. sulfurreducens*. Tools for the genetic manipulation of *G. sulfurreducens* are limited and only the most simple synthetic gene circuits have been adapted for this organism ^33^. The much broader range of strategies for introducing genes and controlling their expression in *E. coli* ^22, 23^ is expected to enable the design and expression of e-PNs with unique properties and functionalities that could not readily be fabricated with *G. sulfurreducens*. Most notably, *E. coli* is an excellent platform for modifying proteins with unnatural amino acids that can confer diverse new functionalities to proteins ^24, 25^. The development of new e-PNs in the *E. coli* chassis for enhanced sensor, electronics, and energy-harvesting applications ^1-8^ is underway.

## Materials and Methods

### *E. coli* strain and culture conditions

*E. coli* NEB 10-beta (New England Biolabs) was grown at 37 °C in LB medium supplemented with appropriate antibiotics as necessary for plasmid preparation, as previously described ^34^. The gene for FimA, the primary monomer for type I pili, was deleted as previously described ^31, 32^ to construct *E. coli* Δ*fimA* (kanamycin sensitive). The strains expressing the modified *E. coli* pilin or the synthetic peptide for assembly into e-PNs were built in this strain.

### Construction of expression vector for type IV pilus assembly

An expression vector for type IV pilus assembly was constructed as described previously ^26^ with several modifications (Figure 1). To construct the basic expression vector for type IV pilus assembly the T7 promoter in the plasmid vector pET24b (Novagen) was replaced with *tac* promoter ^30^. The DNA fragment containing *tac* promoter and *lac* operator was amplified by PCR with a primer pair, Ptac-F/Olac-R (Table S1) and pCD341 ^35^ as template. The PCR product was digested with BglII and XbaI and replaced the BglII-XbaI region containing T7 promoter and *lac* operator in pET24b. The resultant plasmid was designated p24Ptac.

Next, genes for *E. coli* type IV pilus assembly without the major pilin gene were cloned in p24Ptac. The genes include *hofB* (ATPase), *hofC* (platform protein), *hofM* (assembly protein), *hofN* (assembly protein), *hofO* (assembly protein), *hofP* (assembly protein), *hofQ* (secretin), *ppdA* (minor pilin), *ppdB* (minor pilin), *ygdB* (minor pilin), *ppdC* (minor pilin), and *gspO* (prepilin peptidase) ^26^. DNA fragment containing *ppdA, ppdB, ygdB, ppdC*, and *gspO* was prepared by 2-step PCR. Fragments containing *ppdA, ppdB, ygdB*, and *ppdC*, or *gspO* were amplified by PCR with primer pairs, ppdA-F/ppdC-R and gspO-F/gspO-R (Table S1), respectively. The fragment containing *ppdA, ppdB, ygdB, ppdC*, and *gspO* was amplified by PCR with these PCR products as template and a primer pair, ppdA-F/gspO-R. The PCR product was digested with HindIII and XhoI and cloned in pBluescript II SK (Stratagene). The fragment containing *hofM, hofN, hofO, hofP*, and *hofQ* was amplified by PCR with the primer pair, hofM-F/hofQ-R (Table S1). The PCR product was digested with XbaI and HindIII and cloned in the plasmid containing *ppdA, ppdB, ygdB, ppdC*, and *gspO*. The fragment containing *hofB* and *hofC* was amplified by PCR with a primer pair, hofB-F/hofC-R (Table S1). The PCR product was digested with SacI and XbaI and cloned in the plasmid containing *hofM, hofN, hofO, hofP, hofQ, ppdA, ppdB, ygdB, ppdC*, and *gspO*. Fragment containing *hofB, hofC, hofM, hofN, hofO, hofP, hofQ, ppdA, ppdB, ygdB, ppdC*, and *gspO* was prepared by digesting the plasmid containing *hofB, hofC, hofM, hofN, hofO, hofP, hofQ, ppdA, ppdB, ygdB, ppdC*, and *gspO* with SacI and XhoI and cloned in p24Ptac (Figure 1). The resultant plasmid was designated T4PAS/p24Ptac.

### Expression and harvesting of pili comprised of the *E. coli* pilin PpdD

The fragment containing a gene for PpdD with HA tag (Figure 1e) was amplified with a primer pair, ppdD-F/ppdD-HA-R (Table S1), digested with NdeI and SacI, and cloned in T4PAS/p24Ptac. The resultant plasmid was termed ppdD-HA/T4PAS/p24Ptac.

For initial studies with the *E. coli* strain expressing the *E. coli* pilin PpdD, the plasmids ppdD-HA/T4PAS/p24Ptac or T4PAS/p24Ptac were transformed into *E. coli* Δ*fimA*. A single colony from LB agar plate containing kanamycin ^34^ was inoculated in TB medium (Novagen) supplemented with 1% glycerol and kanamycin and incubated at 30° C for 24 h to the stationary phase. Pili were sheared from cells and precipitated with TCA as described previously^26^.

### Expression and harvesting of e-PNs

A fragment encoding a gene for a synthetic pilin monomer, which was similar to the PilA monomer of *G. sulfurreducens* but included the signal sequence of PpdD instead of the original PilA signal sequence (Figure 1f), was amplified with a primer pair, EPS-GspilA-F/GspilA-R (Table S1). The amplified fragment was digested with NdeI and SacI and cloned in T4PAS/p24Ptac. The resultant plasmid, designated GspilA/T4PAS/p24Ptac, was transformed into *E. coli* Δ*fimA*. The resultant strain, designated *E. coli* strain GPN (*<u>G</u>eobacter* <u>p</u>rotein <u>n</u>anowires) was grown on 10 cm diameter culture plates of standard LB medium ^34^ amended with kanamycin, and solidified with agar. After overnight growth at 30 °C, cells were scraped from the surface and suspended in 6 ml of M9 media ^34^. Twenty plates of M9 medium supplemented with 0.5% glycerol, 0.5 mM IPTG, and kanamycin were spread-plated with 300 µl of the suspended cells. The plates were incubated at 30 °C for 48 hours. Cells were harvested from the plates with 1 ml of M9 media (500 µl to scrape, 500 µl to wash) for each plate. The 20 ml suspension of cell scrapings was centrifuged at 4000 rpm for 15 minutes at 4 °C to pellet the cells. The supernatant was discarded and the cells were resuspended in 30 ml of 150 mM ethanolamine (pH 10.5) buffer and poured into a Waring blender. The tubes were washed three times with 20 ml of the ethanolamine buffer, which was also added to the blender. The 90 ml suspension was blended for 2 minutes on low speed. The contents of the blender were transferred to a centrifuge bottle along with a wash of the blender with 10 ml of ethanolamine buffer. The blended material was centrifuged at 5000 x g for 30 minutes at 4°C. The supernatant was collected. Triton X100 detergent was added to provide a final concentration of 6 mM. The mixture was shaken at 100 rpm at room temperature for 45 minutes then added to a stirring filtration unit that had a 100 kDa molecular weight cutoff membrane filter made from polyethersulfone (Omega membrane 100K 76 mm, Pall Corporation). Additional ethanolamine buffer was added to dilute the sample to yield a final Triton X100 concentration of 2 mM. The sample was filtered under nitrogen gas (69 kPa). The sample on the filter was washed four times with 100 ml of water. The e-PNs were collected from the filter by scrapping the surface into 500 µl of water. The scrapping procedure was repeated two more times to yield a suspension of e-PNs in 1.5 ml of water.

### Western blot analysis

The presence of pilin monomers in whole cell extracts and pili preparations was evaluated with Western blot analysis. Whole cell extracts were prepared with B-PER Complete Bacterial Protein Extraction Reagent (Thermo Fisher Scientific). Western blot analysis was conducted as described previously ^7^. PpdD-HA pilin was detected with an anti-HA antibody (HA Tag Polyclonal Antibody, Invitrogen). The *G. sulfurreducens* pilin monomer, PilA, was detected with an anti-PilA antibody ^7^.

### Protein Nanowire Conductance

The conductance of e-PNs expressed in *E. coli* from the *G. sulfurreducens* pilin was analyzed as previously described ^13, 15^. The e-PN preparation in water was adjusted to 500 μg protein per μl. As previously described ^13^, a 2 μl aliquot of the e-PN preparation was dropcast onto the center of three different gold electrode nanoarrays and allowed to dry for 1 hour at 24 °C, after which another 2 μl was dropcast and left to dry overnight at 24 °C. Each of the three electrodes nanoarrays was analyzed with a Keithley 4200 Semiconductor Characterization System setup with four probes to conduct a current-voltage (I-V) curve using a ± 30 × 10^−8^ V sweep with a 5s delay and a 250s hold time. The analyses on each of the three nanoarrays were repeated in triplicate. Thin film conductance was calculated by extracting the slope of the linear fit of the current-voltage response for each of the three measurements on the three electrodes using the formula; G= I/V, where *G* is the conductance, *I* is the current and *V* is the voltage.

In order to evaluate the conductance of individual e-PNs, 100 μl aliquot of a culture of *E. coli* expressing the *G. sulfurreducens* pilin was dropcast onto highly oriented pyrolytic graphite (HOPG) and allowed to sit for 10 minutes. Then the excess liquid was wicked away with a Kimwipe, and an equal volume of deionized water was added to wash off excess salts etc., blotted dry, then dried for 12 hours at 24 °C. Samples were loaded into an Oxford Instruments Cypher ES Environmental AFM and equilibrated for at least 2 hours. The AFM was operated in ORCA electrical mode with a Pt/Ir-coated Arrow-ContPT tip with a 0.2 N/m force constant (NanoWorld AG, Neuchâtel, Switzerland). e-PNs were identified in Amplitude Mapping mode (AM-AFM). Point mode current-voltage spectroscopy was carried out by switching to contact mode and gently touching the conductive tip, which was acting as a translatable top electrode, to the top of the e-PN with a force of 1 nN. A voltage sweep of ± 0.6 V set at 0.99 Hz was applied to three independent points on each of three individual e-PNs. The conductance was calculated, as above, from the slope of the linear fit of the current-voltage response between −0.2 and 0.2 V.

