## Supplemental for "An *Escherichia coli* Chassis for Production of Electrically Conductive Protein Nanowires"

Table S1. Primers used in this study.
Recognition sequences for restriction enzymes are underlined.

| Name | Sequence | Enzyme |
|---|---|---|
| Ptac-F | TTCAGATCTGCAAATATTCTGAAATGAGC | BglII |
| Olac-R | TTCTCTAGAGGGGAATTGTTATCCGCTCAC | XbaI |
| hofB-F | TCTGAGCTCAGGAAGGAGCGGCAATGAATATTC | SacI |
| hofC-R | ATCTCTAGATTATCCCATCCCACTCATC | XbaI |
| hofM-F | ATCTCTAGAAGGCCGTCAGAGTGACGGGTGATAAG | XbaI |
| hofQ-R | TCTAAGCTTACTCACTGGAAACCAGTC | HindIII |
| ppdA-F | TCTAAGCTTAGGAGACTGCCGGCATGAAAACACAAC | HindIII |
| ppdC-R | GTCATTATTGTTGCTCCCTGCTACTGACGATTCGGACAATG | |
| gspO-F | CATTGTCCGAATCGTCAGTAGCAGGGAGCAACAATAATGAC | |
| gspO-R | TCTCTCGAGTTATCTGCAAGCACAGATCC | XhoI |
| ppdD-F | TCTCATATGGACAAGCAACGCGGTTTTAC | NdeI |
| ppdD-HA-R | TCTGAGCTCTTACGCGTAGTCCGGCACGTCGTACGGGTAGTTGGCGTCATCAAAGCGG | SacI |
| EPS-GspilA-F | TCTCATATGGACAAGCAACGCGGTTTCACCCTTATCGAGCTGC | NdeI |
| GspilA-R | TCTGAGCTCTTAACTTTCGGGCGGATAGG | SacI |
